# Precise cortical contributions to feedback sensorimotor control during reactive balance

**DOI:** 10.1101/2023.10.02.560626

**Authors:** Scott Boebinger, Aiden Payne, Giovanni Martino, Kennedy Kerr, Jasmine Mirdamadi, J. Lucas McKay, Michael Borich, Lena Ting

## Abstract

The role of the cortex in shaping automatic whole-body motor behaviors such as walking and balance is poorly understood. Gait and balance are typically mediated through subcortical circuits, with the cortex becoming engaged as needed on an individual basis by task difficulty and complexity. However, we lack a mechanistic understanding of how increased cortical contribution to whole-body movements shapes motor output. Here we use reactive balance recovery as a paradigm to identify relationships between hierarchical control mechanisms and their engagement across balance tasks of increasing difficulty in young adults. We hypothesize that parallel sensorimotor feedback loops engaging subcortical and cortical circuits contribute to balance-correcting muscle activity, and that the involvement of cortical circuits increases with balance challenge. We decomposed balance-correcting muscle activity based on hypothesized subcortically- and cortically-mediated feedback components driven by similar sensory information, but with different loop delays. The initial balance-correcting muscle activity was engaged at all levels of balance difficulty. Its onset latency was consistent with subcortical sensorimotor loops observed in the lower limb. An even later, presumed, cortically-mediated burst of muscle activity became additionally engaged as balance task difficulty increased, at latencies consistent with longer transcortical sensorimotor loops. We further demonstrate that evoked cortical activity in central midline areas measured using electroencephalography (EEG) can be explained by a similar sensory transformation as muscle activity but at a delay consistent with its role in a transcortical loop driving later cortical contributions to balance-correcting muscle activity. These results demonstrate that a neuromechanical model of muscle activity can be used to infer cortical contributions to muscle activity without recording brain activity. Our model may provide a useful framework for evaluating changes in cortical contributions to balance that are associated with falls in older adults and in neurological disorders such as Parkinson’s disease.

## Introduction

Although it is well known that the cortex can fine tune highly automatic behaviors such as walking and balance [1–12], we do not have a mechanistic understanding of how this cortical activity shapes whole body motor output. Neural control of walking and balance is seldom under the exclusive control of either cortical or subcortical processes [13]. Rather, there is a spectrum between cortical and subcortical processes that can shift on an individual basis as difficulty and complexity of the task increases. Models of reactive balance control have previously focused solely on subcortically-mediated muscle activity [14–17]. There is increasing evidence that the cortex becomes engaged during reactive balance control as balance task difficulty increases on an individual basis [11,18–23], but it’s unclear how cortical activity shapes balance-correcting muscle activity [24,25]. To address this gap, we examined perturbation-evoked cortical and muscle activity to investigate how cortical activity relates to motor output. Reactive balance recovery is a robust paradigm to investigate shifts in hierarchical motor control as the task difficulty can easily be manipulated by increasing the magnitude of the balance perturbation.

Cortical contributions to reactive balance control have not been considered in models of balance-correcting muscle activity. The goal of standing balance control is to maintain the body’s center of mass (CoM) in an upright equilibrium over an individual’s base of support [26–28]. Deviations of the CoM from an upright, standing posture can therefore be thought of as balance errors. There is likely to be strong evolutionary pressure to correct these errors as quickly and accurately as possible [29]. Neurophysiological lesion studies have shown that these responses engage spinal and subcortical pathways that enable precise action at the shortest possible latencies [8,9,12,30]. Further, previous computational studies have shown that the patterns of magnitude and timing of balance-correcting muscle responses can be modeled using sensory information encoding balance error via a muscle Sensorimotor Response Model (mSRM) [14–17,31]. In response to a balance perturbation, evoked muscle activity has a characteristic waveform consisting of an initial burst of activity followed by a sustained plateau region of tonic activity [15]. The initial burst, occurring ∼100ms after perturbation onset, is thought to be subcortically-mediated due to its latency and may be akin to the long latency reflex (LLR) observed in both the upper and lower limbs [21,32–34]. Prior work has consistently demonstrated that this initial burst of muscle activity is driven by CoM acceleration at a latency consistent with subcortical processing across individuals, task difficulty, and regardless of age or neurological impairment [15–17]. Later portions of muscle activity (>150ms after perturbation onset) are more variable as muscle activity at this latency may be mediated by both cortical and subcortical sensorimotor circuits [11,21]. Therefore, we added an additional, longer-latency CoM feedback loop to the mSRM, hereby referred to as the hierarchical SRM (hSRM), to account for potential cortical contributions to later phases muscle activity.

Robust cortical signals are evoked following perturbations to standing balance, however it is unclear whether this activity is part of a transcortical sensorimotor feedback loop involved in mediating cortical control of balance-correcting muscle activity. Evoked cortical signals have temporal characteristics similar to perturbation-evoked muscle activity [35–37], but we do not know if these signals are also driven by balance error, quantified as CoM kinematic deviations from upright stance. Recordings from the cortex using electroencephalography (EEG) have revealed a large, negative peak of cortical activity (N1) along with changes in spectral power evoked by perturbations to upright stance [24,38–41]. The cortical N1 is thought of as an error assessment signal evoked when external stimuli cause an unexpected error from the upright posture [36,37,42]. In the time-frequency domain, perturbations to upright stance also evoke an increase in power in the beta frequency range (β; 13-30Hz activity) [43,40,44,24,45,cf. 41]. β activity is involved in sensorimotor processing and motor output, with increased β activity facilitating sensory processing at the expense of movement initiation [46–50]. Taken together, these findings are consistent with the theory that sensorimotor β activity signals the “maintenance of the status quo” where decreases in β activity facilitate changes in an individual’s sensorimotor set while increases indicate favoring the current sensorimotor set [50–53].

Perturbation-evoked cortical N1 and sensorimotor β activity have previously been shown to weakly scale with perturbation magnitude, consistent with increasing balance error [24,25]. However, it has not been quantitatively shown whether feedback of balance error signals can account for evoked cortical responses during balance recovery. Therefore, we adapted the mSRM to reconstruct perturbation-evoked cortical N1 and β activity using the same CoM kinematic error signals, in a model we call the cortical SRM (cSRM). The perturbation-evoked cortical N1 and increase in β activity occur simultaneously with the initial, automatic, subcortically-mediated motor response [35,37] and therefore cannot contribute to this initial burst of muscle activity. However, we do not know if these neurophysiological signatures could contribute to subsequent, longer-latency muscle activity. Therefore, we made two separate adaptations to the hSRM that include either cortical N1 or β activity as a predictor of longer latency balance-correcting muscle activity, rather than CoM kinematics, to test whether these cortical signatures could be involved in a transcortical sensorimotor feedback loop that contributes to the second–but not first–burst of balance-correcting muscle activity.

Here, we hypothesize that parallel sensorimotor feedback loops engaging subcortical and cortical circuits contribute to balance-correcting muscle activity, and that the involvement of cortical circuits increases with balance challenge. We used hierarchical neuromechanical models of reactive balance control to reconstruct cortical and muscle activity evoked by support-surface balance perturbations. Balance perturbations were delivered at varying difficulties to investigate the relationship between sensory information, muscle activity, and cortical activity as balance challenge increases. We predict that when balance task difficulty is sufficiently low, the balance-correcting neuromuscular response is primarily subcortically-mediated (*Figure 1A*) and will therefore be well characterized by the mSRM. We further predict that cortical contributions to balance-correcting muscle activity will appear at higher levels of balance task difficulty and alter the motor response (*Figure 1B*). Additionally, we predict that cortical N1 and sensorimotor β activity are also driven by sensory information encoding balance error and contribute to later phases of balance-correcting muscle activity. Finally, we predict that accounting for potential cortical contributions to motor output will improve reconstruction accuracy of balance-correcting muscle activity regardless of the perturbation difficulty.

**Figure 1:**
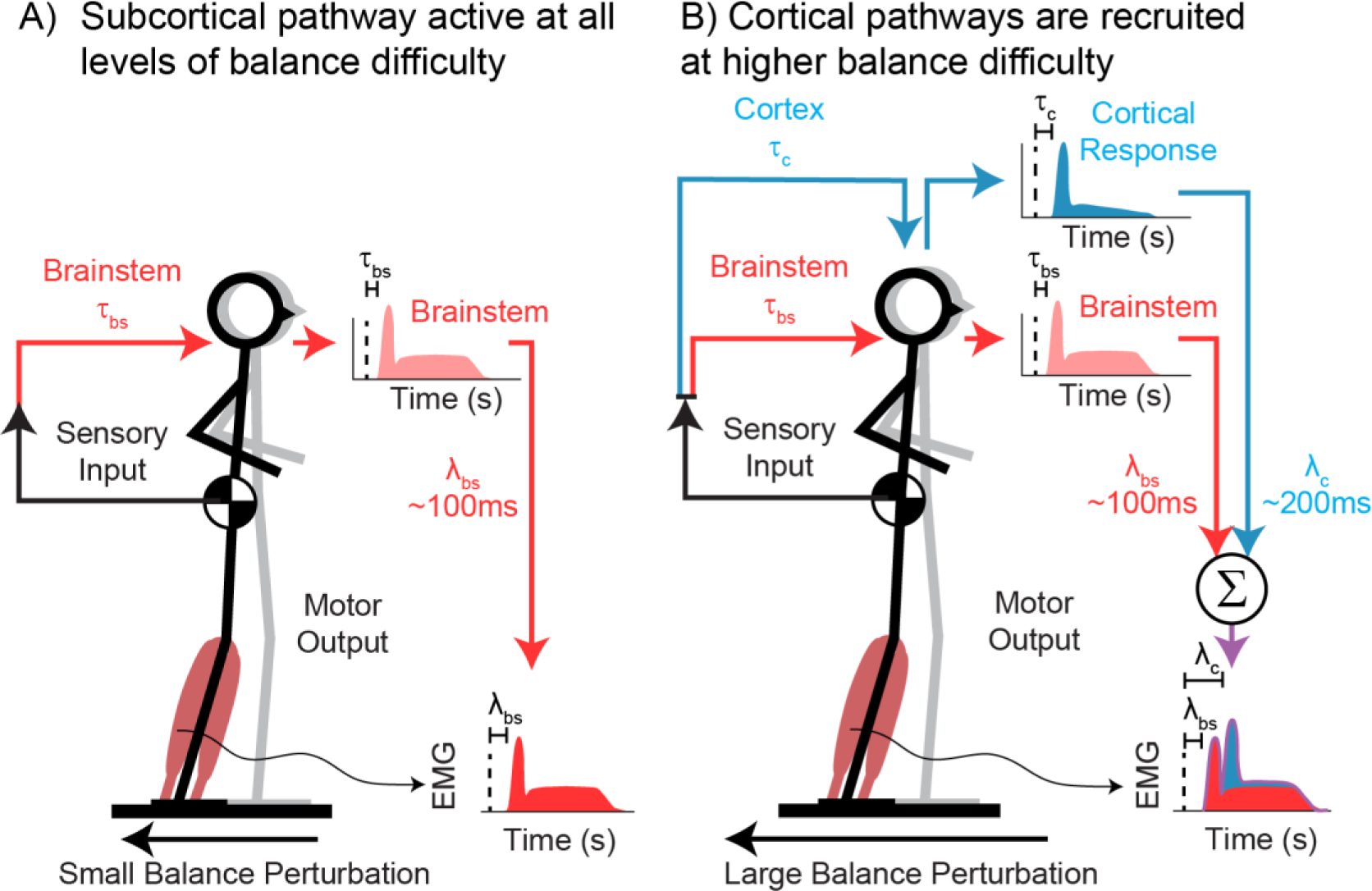
Cortical and subcortical sensory motor loops involved in reactive balance control. A) At low balance difficulty, reactive balance control is primarily mediated through subcortical sensorimotor circuits (red) at an earlier latency (λ_bs_). B) At higher balance difficulty, cortical sensorimotor circuits (blue) contribute to the balance-correcting muscle activity at longer latency (λ_c_).

## Methods

### Participants

Seventeen young adults (10 female, 25 ± 6 years old, 168 ± 8 cm tall, 69 ± 14 kg) were recruited from Emory University and the surrounding Atlanta community to participate in this study. Participants were excluded if they reported having a history of lower extremity joint pain, contractures, major sensory deficits, evidence of orthopedic, muscular, or physical disability, evidence of vestibular, auditory, or proprioceptive impairment, orthostatic hypotension, and/or any neurological insult. All experiments were approved by the Emory University Institutional Review Board. All participants gave informed written consent before participating. Other outcome measures from this cohort have been reported previously [24,25,37].

### Balance perturbations

During the experiment, participants stood barefoot on a motorized platform (Factory Automation Systems, Atlanta, GA, USA). Participants underwent 48 backward translational support-surface perturbations, which were delivered at unpredictable timing and randomized order of magnitudes [24,25,37]. Each participant received an equal number of perturbations in three magnitudes: a small perturbation (7.7 cm, 16.0 cm/s, 0.23 g), which was identical across participants, and two larger magnitudes (medium: 12.6–15.0 cm, 26.6–31.5 cm/s, 0.38–0.45 g, and large: 18.4–21.9 cm, 38.7–42.3 cm/s, 0.54–0.64 g), which were adjusted based on participant height [24,25,37]. To limit predictability, perturbation magnitudes were presented in pseudorandom block orders, with each of the eight blocks containing two perturbations of each magnitude. As previously described [24,25,37], participants were instructed to perform a stepping response in half of the perturbations and to resist stepping in the other half. Only non-stepping trials were included in our analyses because stepping leaves the position of the base of support temporarily undefined, thereby preventing the calculation of kinematic errors between the CoM and the base of support. Successful non-stepping trials were identified using platform-mounted force plates (AMTI OR6-6). To limit fatigue, 5-minute breaks were given every 15 minutes of experimentation.

### Electroencephalography (EEG)

Brain activity was recorded continuously throughout the balance perturbation series using a 32-channel set of actiCAP active recording electrodes (Brain Products GmbH, Munich, Germany) placed according to the international 10–20 system, with the exception of electrodes TP9/10 which were placed directly on the scalp over the mastoids [24].

EEG data were reprocessed with updated routines designed to further reduce muscle and motion artifact using a more rigorous, standardized preprocessing pipeline (*Figure 2*), as the previous analysis may not have adequately reduced muscle and motion artifact. Data were pre-processed in Matlab 2022b (Mathworks, Natick, MA, USA) using custom scripts and EEGLAB [54] functions that down sampled from 1000Hz to 500Hz, applied a 1Hz high-pass filter, removed and interpolated bad channels, minimized line noise [55,56], and applied an average referencing method. Data were epoched -500 to 2000ms relative to perturbation onset, and decomposed into maximally independent components for non-neural artifact removal [57].

**Figure 2:**
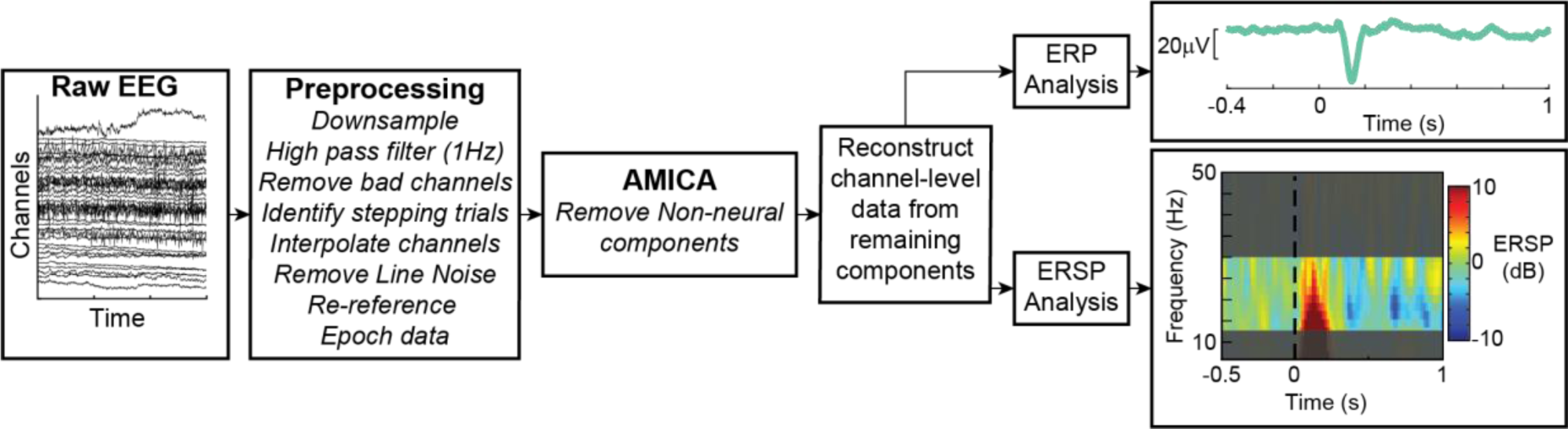
Electroencephalography (EEG) processing pipeline. Raw EEG is preprocessed, non-neural artifacts are removed, and remaining brain components are further analyzed in electrode space to quantify perturbation evoked cortical N1 and sensorimotor β activity.

**Figure 3:**
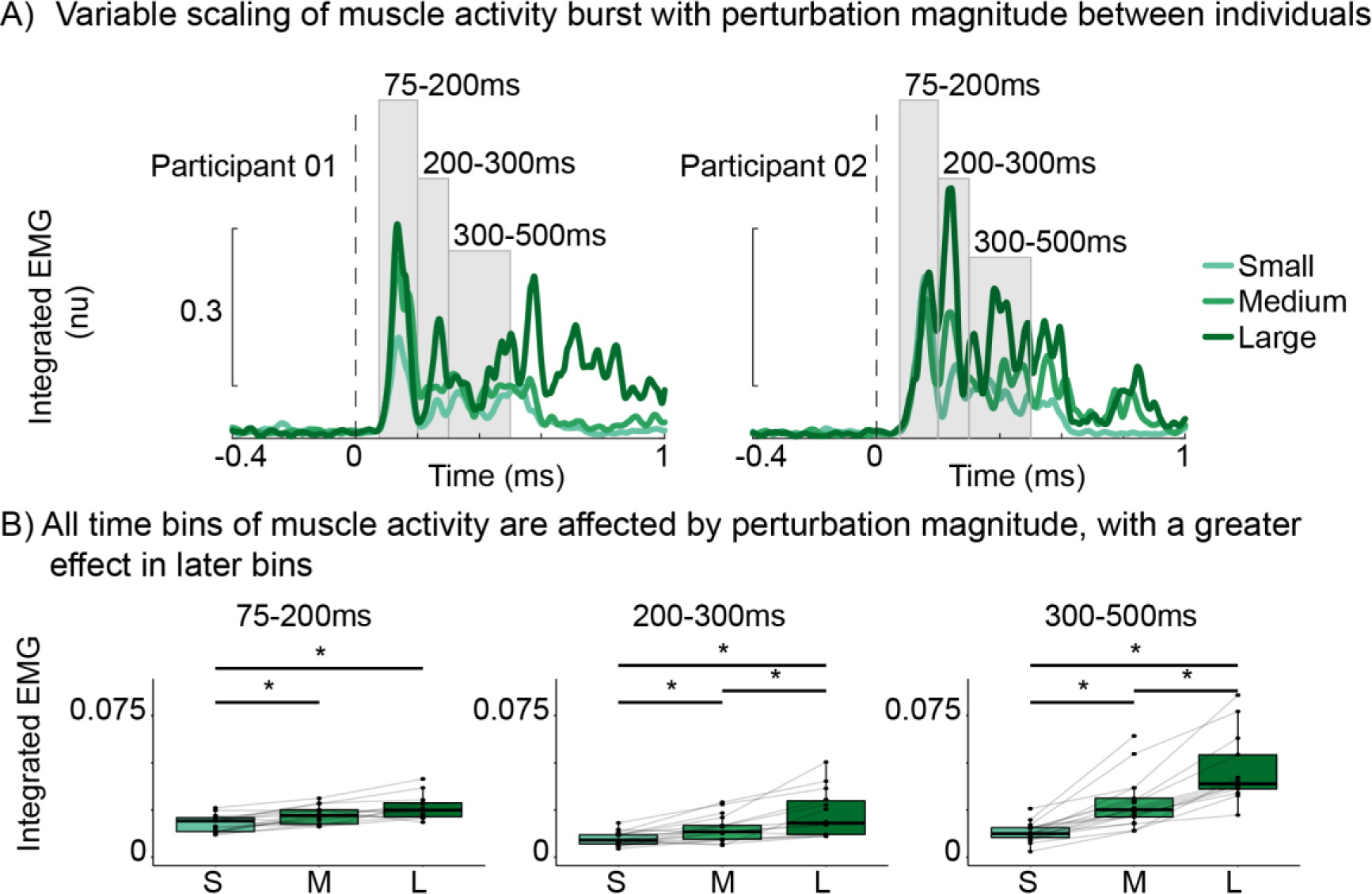
Perturbation-evoked muscle activity increases as balance challenge increases. A) Exemplar participant at all three perturbation magnitudes: small (light green), medium (green), and large (dark green). B) group averaged data for each time bin. *p<0.05.

After removal of non-neural components, channel-level data were reconstructed from the remaining components, and analyses focused on the Cz electrode positioned over primary motor and supplementary motor cortical regions. N1 amplitude was calculated as the minimum value for the event related potential 100-200ms post-perturbation relative to a baseline window (400ms to 100ms pre-perturbation). Time–frequency analyses were performed on preprocessed EEG data to assess β power prior to and during reactive balance recovery. Changes in spectral power were quantified relative to a baseline window (400ms to 100ms pre-perturbation) in single-trial epochs using wavelet time–frequency analyses in EEGLAB (pop_newtimef.m). A tapered Morlet wavelet with three cycles at the lowest frequency (6Hz), linearly increasing up to 23 cycles at the highest frequency (50Hz) was used to measure power at each frequency in a sliding window of 256ms. This wavelet transformation calculates the event-related spectral perturbation (ERSP), which represents changes in power relative to perturbation onset in a defined set of frequencies [58]. Single-trial ERSP values were averaged across sampled frequencies within the β range (13-30Hz) to obtain a single waveform of the time course of β power for each trial. Single-trial β power values were then averaged across trials within each perturbation condition for each participant. Condition averaged β power was then separated into three separate time bins (50-150ms, 150-250ms, and 250-500ms). These time bins were selected to allow for comparison with previous studies on this same dataset [24].

### Electromyography (EMG)

Surface EMGs (Motion Analysis Systems, Baton Rouge, LA) were collected bilaterally from the tibialis anterior (TA), medial gastrocnemius muscle (MG), and sternocleidomastoid (SC) muscles bilaterally. Analysis focused on the MG since this muscle is the primary agonist for a backward perturbation of the support surface. Skin was shaved if necessary and scrubbed with an isopropyl alcohol wipe before electrode placement using standard procedures [59]. Bipolar silver silver-chloride electrodes were used (Norotrode 20, Myotronics, Inc., Kent, WA, USA). Electromyography signals were sampled at 1000 Hz and anti-alias filtered with an online 500 Hz low-pass filter. Raw EMG signals were epoched between -400ms and 1400ms relative to perturbation onset. EMG signals were then high-pass filtered at 35Hz offline with a sixth-order zero-lag Butterworth filter, mean-subtracted, half-wave rectified, and subsequently low-pass filtered at 40Hz [16,17,24,37]. Single-trial EMG data were normalized to a maximum value of 1 across all trials within each participant for left and right sides independently. EMG data were then averaged across trials within each perturbation magnitude for each participant. Perturbation evoked muscle activity was then separated into three separate time bins (75-200ms, 200-300ms, and 300-500ms). These time bins were selected based on visual inspection of all participants’ data to ensure that they captured the initial burst of muscle activity (75-200ms) as well as the longer latency bursts (200-300ms and 300-500ms).

### Statistical characterization of EMG and EEG responses

Statistical tests were performed in RStudio version 1.4.1717 (R Core Team, Vienna, Austria). Differences in cortical N1, sensorimotor β activity, and muscle activity between perturbation magnitudes were assessed using two-way ANOVAs with perturbation magnitude as a group factor and participant included as a random factor. Separate ANOVAs were performed for each prespecified time bin. Post hoc comparisons were performed by comparing the estimated marginal means of these prespecified time bins between perturbation magnitudes. Tests were considered statistically significant at p ≤ 0.05.

### Sensorimotor Response Models (SRMs)

To investigate the relationship between sensory information, cortical activity, and muscle activity, we used a series of delayed feedback control models to reproduce neurophysiological activation patterns based on sensory predictors.

### Muscle Sensorimotor Response Model (mSRM)

First, we used the classic mSRM to reconstruct balance-correcting muscle activity using a single feedback loop (*Figure 4A*). We assume the mSRM reconstructs subcortically-mediated muscle activity due to the short latency of the transformation between balance errors and the muscle activity that is reconstructed by this model. The mSRM reconstructs balance-correcting muscle activity using kinematic signals of balance error [17]. CoM displacement (d), velocity (v), and acceleration (a) relative to the base of support were each weighted by a feedback gain (k_d1_, k_v1_, k_a1_), summed, and delayed be a common time delay (λ_bs_) to account for ascending and descending neural transmission and processing in subcortical circuits:

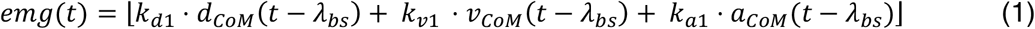

**Figure 4:**
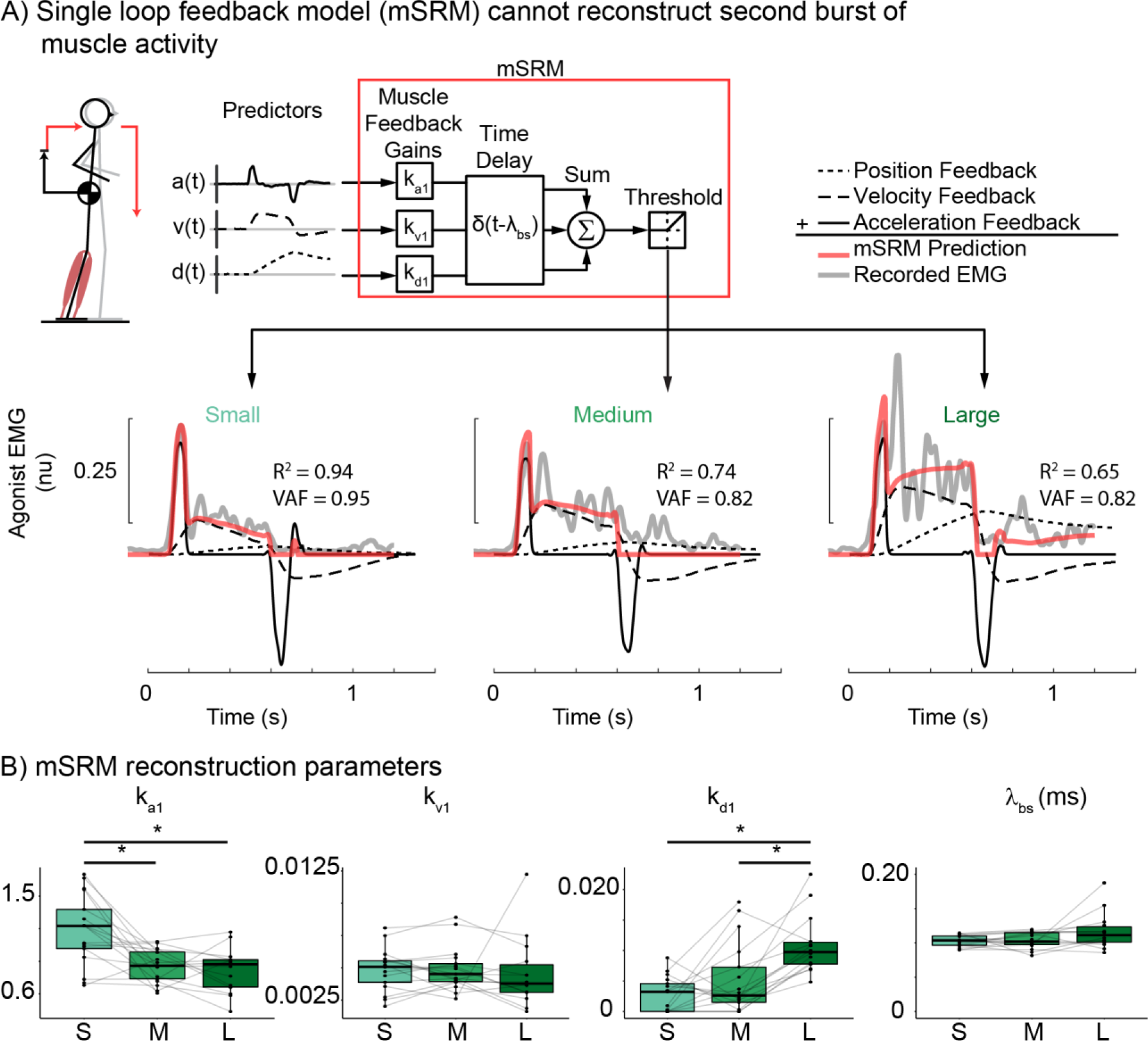
Single loop muscle sensorimotor response model (mSRM) A) mSRM schematic and reconstruction of exemplar participant data (Participant 02) at each perturbation magnitude. mSRM reconstructs balance-correcting muscle activity as the weighted sum of delayed center of mass kinematics. B) group mSRM parameters for CoM acceleration (k_a1_), velocity (k_v1_), and displacement (k_d1_) gains as well as time delay (λ_bs_). *p<0.05

The reconstructed signal (emg(t)) is half wave rectified to represent excitatory drive motor pools [17].

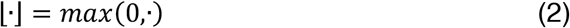

### Hierarchical sensorimotor response model (hSRM)

Here we augmented the preexisting mSRM to include hierarchical feedback loops to account for potential cortical contributions to balance-correcting muscle activity (hSRM, *Figure 5A*). The hSRM reconstructs subcortically-mediated balance-correcting muscle activity in the same way as the mSRM. However, the hSRM has an additional predictor of cortical contributions (f_cortex_(t)) to balance-correcting muscle activity at a longer latency (*Equation 3*).

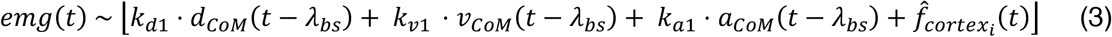

**Figure 5:**
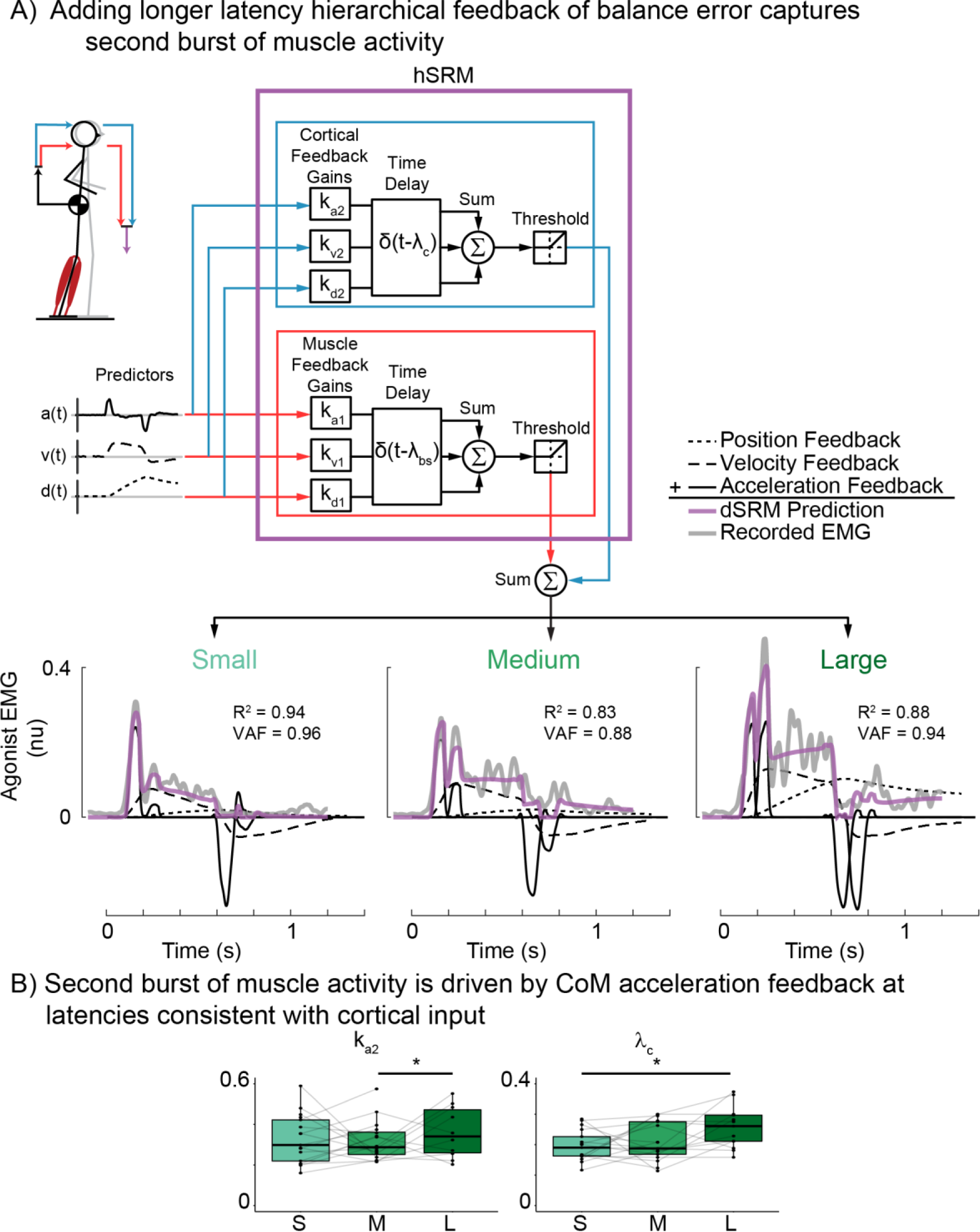
Double loop hierarchical sensorimotor response model (hSRM) A) hSRM schematic and reconstruction of exemplar participant data (Participant 02) at each perturbation magnitude. hSRM adds an additional, longer-latency, cortical feedback loop (blue lines) to the single-latency mSRM (red lines) B) group hSRM parameters for the cortical feedback loop CoM acceleration (k_a2_) and time delay (λ_c_). *p<0.05

**Figure 6:**
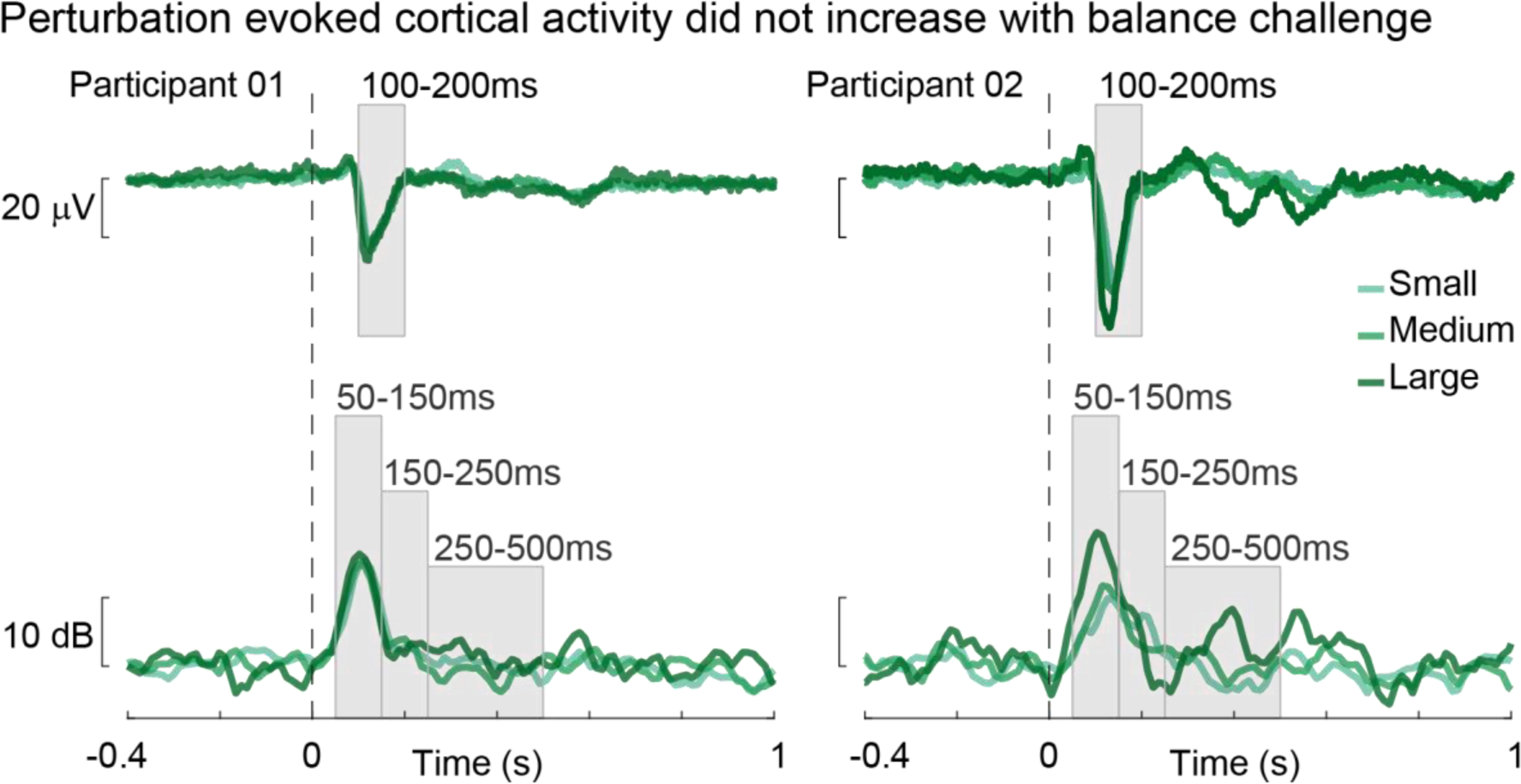
Perturbation-evoked cortical activity as balance challenge increases. Exemplar participants at all three perturbation magnitudes: small (light green), medium (green), and large (dark green).

We created three separate hSRMs based on hypothesized ways the cortex could drive later phases of muscle activity. The first of these models does not incorporate measured cortical activity but supposes that the same sensorimotor transformation from balance error to muscle activity also occurs at the cortical level at a longer latency (λ_c_) than the subcortical circuit (*Figure 5A; Equation 4*).

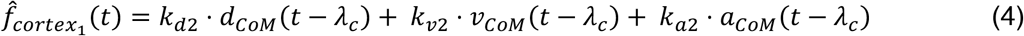

We assume the cortical contribution to be independent of the subcortical process, and therefore weighted CoM kinematic error signals by additional feedback gains (k_d2_, k_v2_, k_a2_) that are independent of those used in the subcortical model (*Equation 1*).

The second of these models incorporates the measured cortical N1 timeseries as a predictor of later phases of muscle activity (*Figure 8A, Equation 5*):

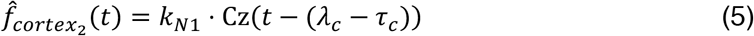

**Figure 7:**
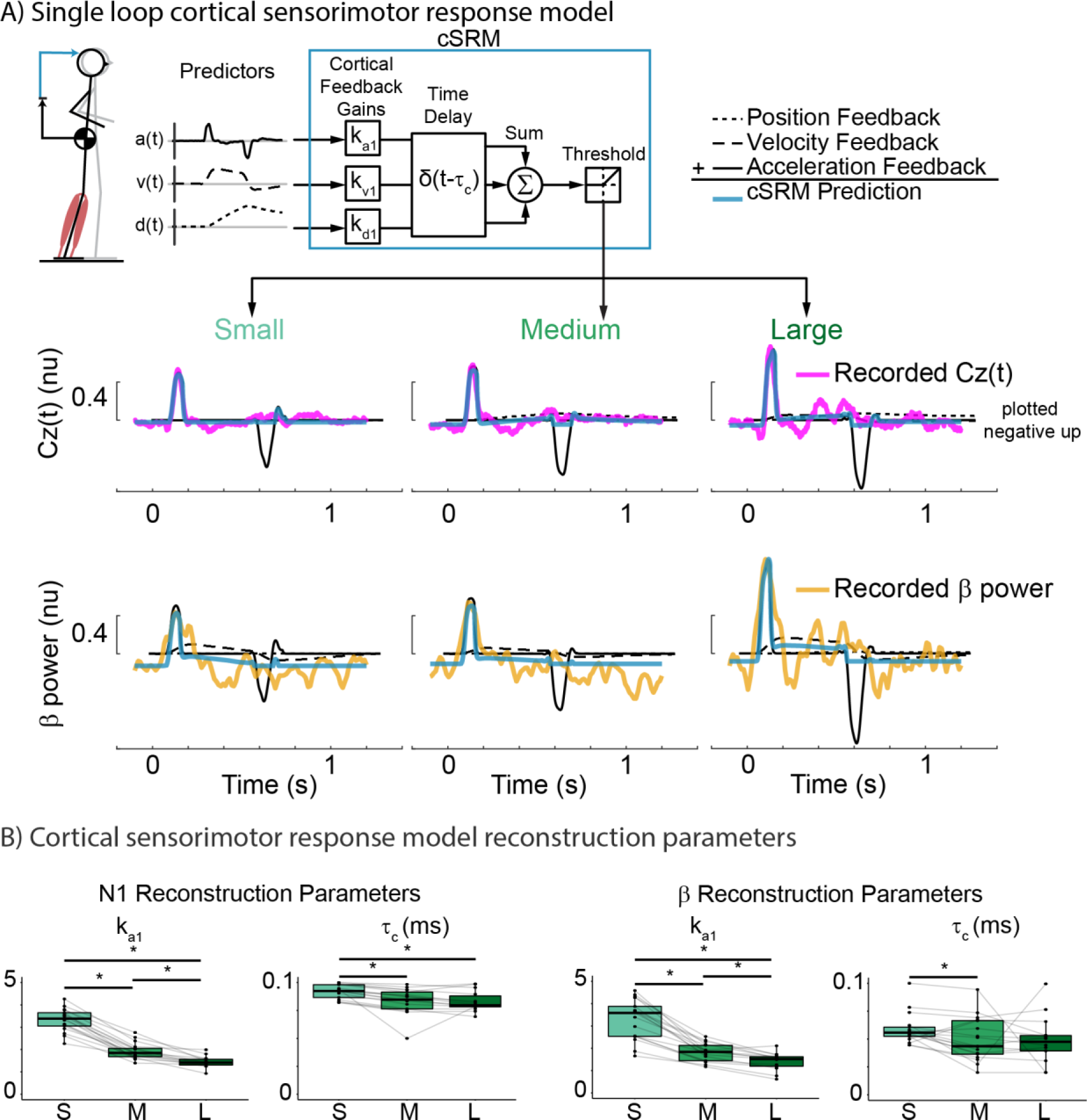
Single loop cortical sensorimotor response model (cSRM) A) cSRM schematic and reconstruction of exemplar participant EEG data (Participant 02) at each perturbation magnitude. cSRM reconstructs either perturbation-evoked cortical N1 (plotted negative up) or sensorimotor β activity as the weighted sum of delayed CoM kinematics. B) group cSRM parameters for CoM acceleration feedback gain (k_a1_) and the associated time delay (τ_c_). *p<0.05

**Figure 8:**
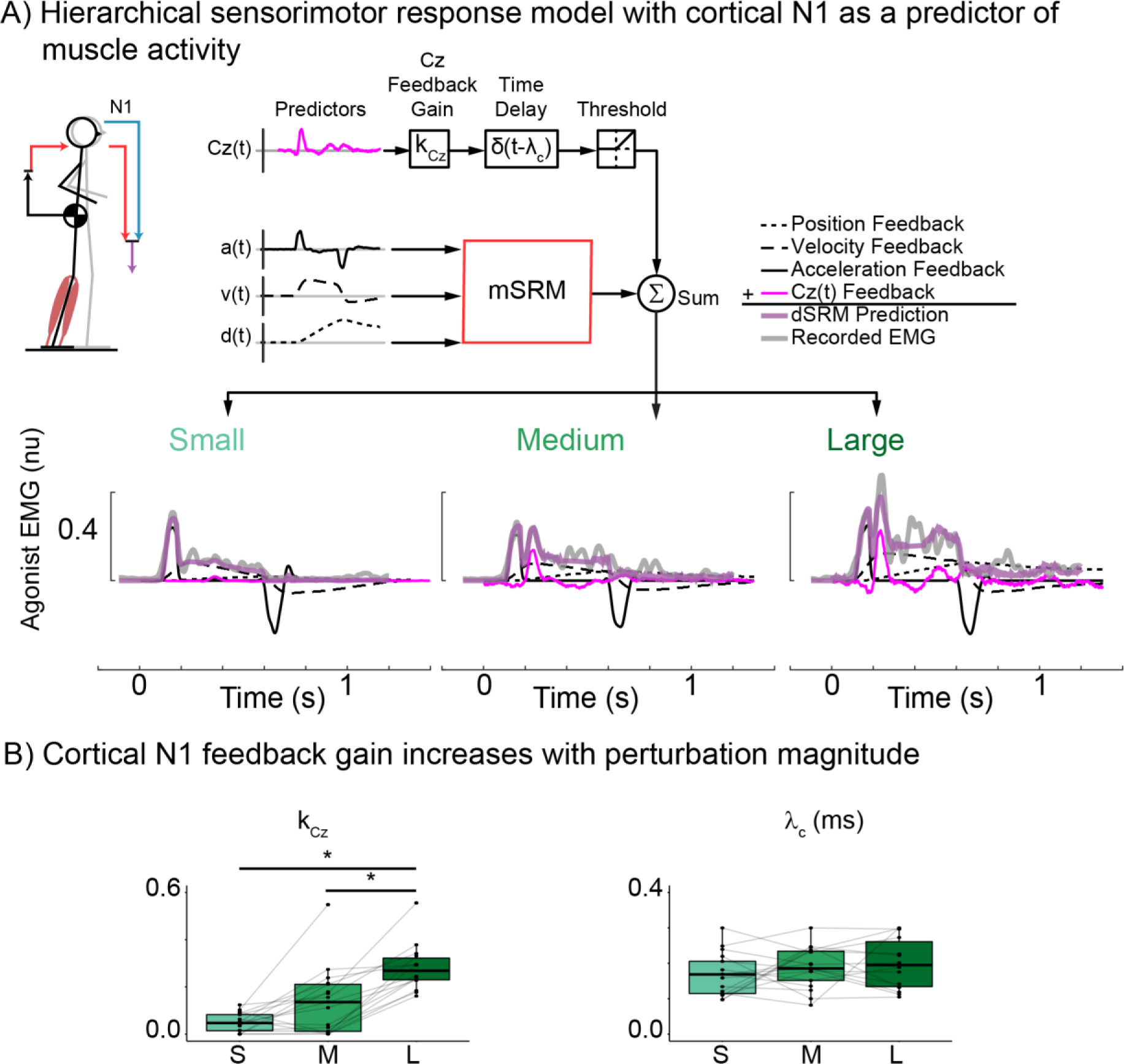
Double loop hierarchical sensorimotor response model (hSRM) using recorded N1 as a predictor of muscle activity. A) hSRM schematic and reconstruction of exemplar participant data (Participant 02) at each perturbation magnitude. hSRM adds an additional, longer-latency, cortical feedback loop (blue lines) to the single-latency mSRM (red lines) B) group hSRM parameters for the cortical feedback loop using N1 (k_Cz_) as a predictor of muscle activity. *p<0.05.

The third of these models includes the measured β activity time series as a predictor of later phases of muscle activity (*Figure 9A, Equation 6*):

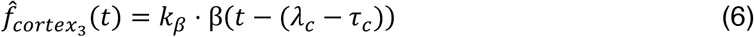

**Figure 9:**
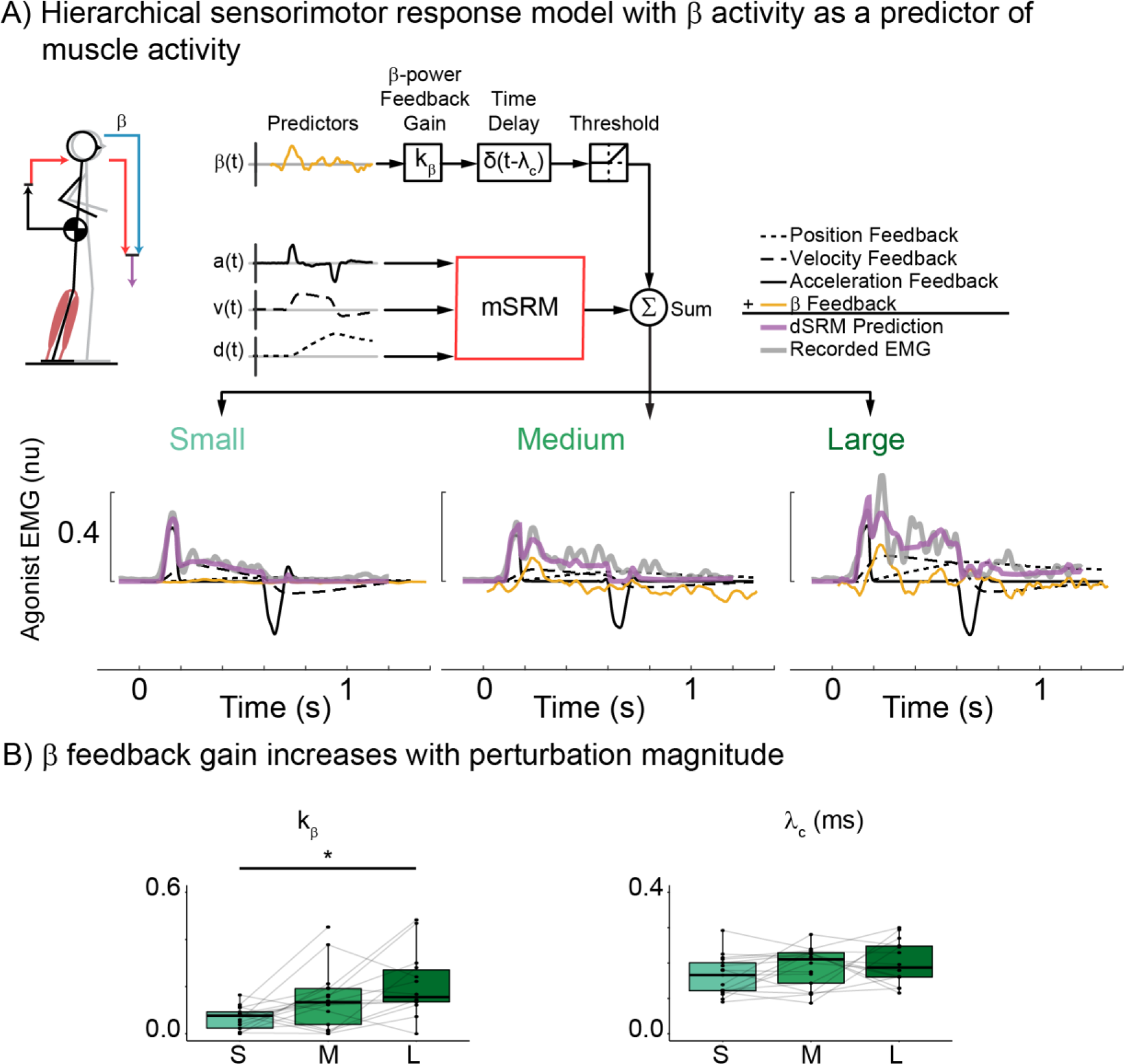
Double loop hierarchical sensorimotor response model (hSRM) using recorded β activity as a predictor of muscle activity. A) hSRM schematic and reconstruction of exemplar participant data (Participant 02) at each perturbation magnitude. B) group hSRM parameters for the cortical feedback loop using β (k_β_) as a predictor of muscle activity. *p<0.05.

Where N1 and β activity were both weighted by independent feedback gains (k_N1_ and k_β_, respectively), and τ_c_ is the latency for the ascending sensory information to reach the cortex.

### Cortical sensorimotor response model (cSRM)

Additionally, we modified the mSRM to reconstruct cortical activity (cSRM) to determine if cortical N1 and sensorimotor β activity could also be driven by sensory information encoding balance error. The cSRM predicts cortical N1 and sensorimotor β activity using kinematic signals of balance error, weighted by feedback gains (k_d1_, k_v1_, k_a1_) and delayed by a shortened time delay (τ_c_) relative to the mSRM to account for the time required for only ascending neural transmission and processing (*Figure 7A; Equation 7 & 8*):

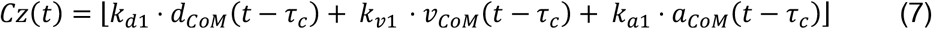

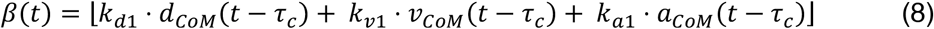

The cSRM reconstructs Cz(t) and β(t) separately, therefore each cSRM reconstruction has an independent set of feedback gains as we assume these two neurophysiological responses are not identical to one another.

### Model parameter identification

For all SRMs mentioned above, model parameters were selected by minimizing the error between recorded neurophysiological data and SRM reconstruction. The reconstruction error term was quantified as the sum of squared reconstruction errors at each time sample as well as the maximum observed reconstruction error (*Equation 9*):

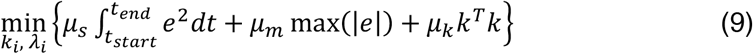

The first term penalizes the squared error (e^2^) between the condition-averaged neurophysiological activity and the SRM reconstruction weighted by μ_s_. The second term penalizes the maximum error between the condition-averaged neurophysiological activity and the SRM reconstruction with a weight μ_m_. The third term penalizes the magnitude of the gain parameters (k_i_) with weight μ_k_ in order to improve convergence when feedback channels do not contribute to the SRM reconstruction. The ratio of weights μ_s_:μ_m_:μ_k_ was 1:1:1e-6. All optimizations were performed in Matlab 2022b (Mathworks, Natick, MA, USA) using the interior-point algorithm in *fmincon*.*m*, as described previously in literature [17]. For single looped models (mSRM and cSRMs) a single optimization was performed to identify model specific feedback parameters (k_d1_, k_v1_, k_a1_).

For double looped models (hSRMs), a separate optimization was performed to identify the shorter latency (k_d1_, k_v1_, k_a1_, λ_bs_) and longer latency (k_d2_, k_v2_, k_a2_, λ_c_) model parameters. The optimization for the shorter latency model parameters is the same as that described for the single loop models. The optimization for the longer latency parameters was performed by reconstructing the residual neurophysiological data, quantified as the difference between the recorded and reconstructed neurophysiological data. After these two separate optimizations were performed, the shorter latency (k_d1_, k_v1_, k_a1_, λ_bs_) and longer latency (k_d2_, k_v2_, k_a2_, λ_c_) model parameters were concatenated into an initial guess for final optimization to allow for modifications as the shorter latency model parameters may have been exaggerated in order to fit longer latency muscle activity. Lower and upper bounds for the gain parameters were ±10% of the initial guess values, lower and upper bounds for the delay parameters were ±10ms of the initial guess values. In all cases, additional parameters supplied to *fmincon*.*m* were as follows: *TolX*, 1e-9; *MaxFunEvals*, 1e5; *TolFun*, 1e-7, with remaining parameters set to default [17].

### Goodness of fit

The goodness of fit between SRM reconstructions and recorded neurophysiological data was assessed using a coefficient of determination (R^2^) as well as variability accounted for (VAF). R^2^ was calculated using the built-in function *regress*.*m* in Matlab 2022b (Mathworks, Natick, MA, USA), and VAF was defined as 100*the square of Pearson’s uncentered correlation coefficient, as performed in previous studies [17,60]. Differences in R^2^ and VAF between perturbation magnitudes were assessed using two-way ANOVAs with perturbation magnitude as a group factor and participant included as a random factor. Tests were considered statistically significant at p ≤ 0.05.

### Model selection

Model selection was performed using Akaike’s Information Criterion (AIC; Equation 10).

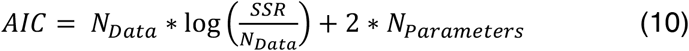

Where N_Data_ is the number of data points reconstructed by the SRMs, SSR is the sum squared error of the SRM reconstruction, and N_Parameters_ is the number of parameters included in the SRM. AIC values were calculated for each SRM described above. An AIC comparison between the mSRM and hSRM was performed to determine whether the addition of a longer latency loop improves the model fit to the data. Differences in AIC between models ≥ 2 were considered meaningful.

A separate AIC analysis was performed to determine if perturbation-evoked cortical signatures are driven primarily by CoM acceleration feedback by comparing cSRM reconstructions when including all the CoM kinematics as predictors or using only CoM acceleration as a predictor. Similarly, we also assessed whether the longer latency burst of muscle activity reconstructed is primarily driven by CoM acceleration feedback by comparing AIC values when using all CoM kinematics, or just CoM acceleration as predictors of cortical contributions to balance-correcting muscle activity in the hSRM.

Additionally, comparisons of reconstruction accuracy between SRMs were performed using a mixed linear effects model with an interaction between the model type (i.e., mSRM or hSRM) and perturbation magnitude with participant included as a random factor. Post hoc comparisons were then performed by comparing the estimated marginal means between the prespecified model types.

## Results

### Variable scaling of muscle activity with perturbation magnitude between participants

During reactive balance recovery, there were individual differences in the response of the first and second burst of muscle activity to increased perturbation magnitude (*Figure 3A*). For example, participant 01’s initial burst (75-200ms) of muscle activity increased between the small and medium perturbations while their second burst (200-300ms) of muscle activity was relatively small or absent. Interestingly, at the large perturbation, the initial burst did not increase further, but a second burst emerged (*Figure 3A*, left panel). Conversely, participant 02 had both the first and second burst already evident in the small perturbation and the first burst remained less affected by perturbation magnitude while the second burst increased markedly (*Figure 3A*, right panel).

At the group level, we investigated the scaling of muscle activity by quantifying the integrated EMG activity within separate time bins that captured the initial and second burst. Across participants, we found a significant increase in integrated EMG across perturbation magnitude for the initial 75-200ms time bin (F(1.54, 19.96) = 21.57, p <0.001), 200-300ms time bin (F(1.31, 17.06) = 17.16, p < 0.001), and 300-500ms time bin (F(1.24, 16.06) = 39.72, p < 0.001) (*Figure 3B*). Post hoc comparisons between perturbation magnitudes showed that in the initial 75-200ms time bin integrated EMG was significantly increased from the small perturbation compared to the medium and large perturbation magnitudes (p < 0.01 for both), while it did not differ between the medium and large perturbation (p = 0.07). In the 200-300ms and 300-500ms time bins, integrated EMG increased across all comparisons of perturbation magnitudes (p < 0.01 for all).

### Single-delay feedback model of balance error fails to capture second burst of muscle activity

At the small perturbation magnitude, balance-correcting muscle activity adheres to the characteristic waveform of an initial burst driven primarily by CoM acceleration error feedback followed by a plateau region driven by CoM velocity and displacement error feedback [14–16]. Therefore, balance-correcting muscle activity is well described by the mSRM at this small perturbation magnitude (*Figure 4A*, small perturbation reconstruction), (R^2^ = 0.82±0.014, VAF = 0.85±0.010). However, as the perturbation magnitude increases, a longer latency burst of muscle activity begins to appear that cannot be described by the single-delay feedback model (*Figure 4A*, gray trace, large perturbation reconstruction) (R^2^ = 0.60±0.038, VAF = 0.81±0.037). mSRM reconstruction accuracy was high in the smallest perturbation (R^2^ = 0.82±0.014, VAF = 0.85±0.010), but worsened in the medium (R^2^ = 0.77±0.014, VAF = 0.85±0.006) and large (R^2^ = 0.60±0.038, VAF = 0.81±0.047) perturbation magnitudes. We found a decrease in mSRM reconstruction accuracy in both R^2^ and VAF as perturbation magnitude increased (R^2^: F(1.86, 26.08) = 32.70, p < 0.001; VAF: F(1.74, 24.41) = 4.37, p = 0.028).

### Adding hierarchical feedback of balance error reconstructs second burst of muscle activity

Accounting for longer latency, potentially cortical contributions to balance-correcting muscle activity enabled reconstruction of the second burst of muscle activity that increases with perturbation magnitude (*Figure 5*). We tested whether the same sensorimotor transformation from balance error to balance-correcting muscle activity occurs at both subcortical and cortical levels, by adding an additional, longer-latency, hierarchical feedback loop (*Figure 5A* blue lines) to the to the mSRM (*Figure 5A* red lines). hSRM reconstruction accuracy was highest in the smallest perturbation (R^2^ = 0.86±0.012, VAF = 0.89±0.008) and medium perturbation (R^2^ = 0.84±0.011, VAF = 0.91±0.005), but decreased slightly in the largest perturbation magnitude (R^2^ = 0.70±0.042, VAF = 0.87±0.050). We found a decrease in hSRM reconstruction accuracy in both R^2^ and VAF as perturbation magnitude increased (R^2^: F(1.47, 20.53) = 17.48, p < 0.001; VAF: F(1.79, 25.09) = 5.76, p = 0.011).

The inclusion of the longer latency feedback loop in the hSRM resulted in more accurate reconstructions of muscle activity across all perturbation magnitudes compared to the single-loop mSRM (R^2^: t(74) = -4.678, p <0.0001; VAF: t(74) = -6.336, p <0.0001). To further compare these models, we performed an AIC analysis and found that the hSRM (AIC = -366379) was a better model of perturbation evoked muscle activity than the single-loop mSRM (AIC = -352106).

A separate AIC analysis was performed to assess whether longer-latency muscle activity was primarily driven by CoM acceleration feedback. We found that the hSRM that only included CoM acceleration feedback (AIC = -383413) was a more appropriate model than the hSRM that included all CoM kinematic feedback (AIC = -366379). Therefore, the second burst was captured by the longer-latency acceleration feedback error. k_a2_, which defines the input-output relationship between CoM acceleration and the second burst of muscle activity, increased with perturbation magnitude (F(1.62, 21.01) = 5.75, p = 0.014) as did the associated time delay (λ_c_) (F(1.86, 24.22) = 4.91, p = 0.018) (*Figure 5B*).

### Perturbation-evoked cortical activity did not increase with balance challenge

In the time-domain, recordings from central midline electrode (Cz) show our perturbation paradigm was able to robustly evoke an N1 response in each participant (*Figure 6*). Across participants, N1 had an average amplitude of -30.6±1.7μV and was observed at 137±1.6ms. Contrary to previous findings in the same data set, N1 amplitude did not significantly increase with perturbation magnitude at the group level (F(1.36, 17.62) = 2.03, p = 0.17). Post hoc comparisons between perturbation magnitude showed N1 amplitude did not differ between any perturbation magnitude (p > 0.25 for all). However, in corroboration with previous findings, N1 latency decreased as perturbation magnitude increased (F(1.22, 19.47) = 10.85, p = 0.002). Post hoc comparisons between perturbation magnitudes showed that N1 latency was significantly increased from the small perturbation compared to the medium and large perturbation magnitudes (p < 0.01 for both), while it did not differ between the medium and large perturbation (p = 0.10).

In the frequency domain, β power increased after perturbation onset and reached a maximum around the latency of the cortical N1. Contrary to what was found previously, this initial rise in β power 50-150ms time bin did not significantly increase with perturbation magnitude at the group level (F(1.86, 25.98) = 2.87, p = 0.078). β power the 150-250ms time bin did not have a significant increase with perturbation magnitude (F(1.98, 27.73) = 2.03, p = 0.15) while the 250-500ms time bin did (F(1.99, 27.89) = 6.37, p = 0.005). Post hoc comparisons between perturbation magnitudes showed that in the 50-150ms and 150-250ms time bins β power was did not significantly increase between any perturbation magnitude (p > 0.05 for all). In the 250-500ms time bin, β power did not significantly increase between the small and medium perturbation magnitude (p = 0.60) nor small and large (p = 0.06) but did significantly increase between the medium and large perturbation magnitude (p = 0.01).

### Single-delay feedback model of balance error can reconstruct perturbation-evoked cortical activity

We tested whether perturbation-evoked cortical N1 and β activity could be explained by sensorimotor transformation of balance error signals using the cSRM (*Figure 7A*). The cSRM was able to reconstruct both cortical N1 and sensorimotor β activity, suggesting that these cortical signatures may also be driven by sensory information encoding balance error. cSRM reconstruction accuracy of the cortical N1 was highest in the smallest perturbation (R^2^ = 0.75±0.12, VAF = 0.78±0.011) with a slight decrease in the medium (R^2^ = 0.72±0.21, VAF = 0.77±0.015) and largest (R^2^ = 0.69±0.041, VAF = 0.77±0.046) perturbation magnitudes. However, this decrease in reconstruction accuracy with perturbation magnitude did not reach significance (R^2^: F(1.70, 23.75) = 1.12, p = 0.33; VAF: F(1.55, 21.68) = 0.10, p = 0.86).

An AIC analysis was performed to assess whether perturbation evoked cortical N1 activity was primarily driven by CoM acceleration feedback. We found that the cSRM that only included CoM acceleration feedback (AIC = -236607) was a more appropriate model than the cSRM that included all CoM kinematic feedback (AIC = -188573). The cSRM acceleration feedback gain, k_a1_, which defines the input-output relationship between CoM acceleration and the cortical N1, decreased with perturbation magnitude (F(1.51, 19.62) = 158.81, p < 0.01) as did the associated time delay (τ_c_) (F(1.31, 17.05) = 10.08, p = 0.003).

cSRM reconstruction accuracy of sensorimotor β activity was lower overall than that of the cortical N1 for all perturbation magnitudes. cSRM reconstruction accuracy of sensorimotor β activity was highest in the smallest (R^2^ = 0.55 ±0.030, VAF = 0.54±0.036) and medium (R^2^ = 0.55±0.021, VAF = 0.50±0.030) perturbation magnitude, but decreased in the largest (R^2^ = 0.43±0.35, VAF = 0.49±0.041) perturbation magnitudes. There was a significant effect of perturbation magnitude on cSRM R^2^, but not VAF, when reconstructing sensorimotor β activity (R^2^: F(1.42, 19.88) = 7.70, p = 0.007; VAF: F(1.97, 27.61) = 1.67, p = 0.207).

We found that the cSRM that only included CoM acceleration feedback (AIC = -184883) was a more appropriate model than the cSRM that included all CoM kinematic feedback (AIC = -177989). The cSRM acceleration feedback gain, k_a1_, which defines the input-output relationship between CoM acceleration and the sensorimotor β activity, decreased with perturbation magnitude (F(1.20, 15.64) = 73.48, p < 0.001) while there was no significant effect with the associated time delay (τ_c_) (F(1.32, 17.10) = 2.11, p = 0.161) (*Figure 7B*).

### Adding hierarchical feedback of perturbation evoked cortical activity reconstructs second burst of muscle activity

To further test whether cortical N1 and sensorimotor β activity could explain later phases of muscle activity we modified the double looped hSRM model to include either N1 (*Figure 8*) or sensorimotor β activity (*Figure 9*) instead of the delayed secondary sensorimotor transformation.

When using N1 as a predictor of muscle activity, reconstruction accuracy was highest in the smallest perturbation (R^2^ = 0.84 ±0.013, VAF = 0.87±0.009) and medium perturbation (R^2^ = 0.82±0.013, VAF = 0.89±0.006), but decreased in the largest perturbation magnitude (R^2^ = 0.67±0.041, VAF = 0.85±0.049). The inclusion of cortical N1 as a predictor of balance-correcting muscle activity allowed the hSRM to capture the notable second burst of balance-correcting muscle activity (*Figure 8A*) that is observed as balance challenge increases. The inclusion of the longer latency feedback loop in the hSRM resulted in more accurate reconstructions of muscle activity across all perturbation magnitudes compared to the single-loop mSRM (R^2^: t(74) = -3.301, p = 0.0015; VAF: t(74) = -3.901, p = 0.002). To further compare these models, we performed an AIC analysis and found that the hSRM when using cortical N1 as a predictor (AIC = -363742) was a better model of perturbation evoked muscle activity than the single-loop mSRM (AIC = -352106). The cSRM feedback gain associated with N1’s contribution to balance-correcting muscle activity (k_Cz_) significantly increased with perturbation magnitude (F(1.96, 25.49) = 46.75, p < 0.001) (*Figure 8B*).

When using sensorimotor β activity as a predictor of muscle activity, reconstruction accuracy was highest in the smallest perturbation (R^2^ = 0.85±0.012, VAF = 0.87±0.008) and medium perturbation magnitudes (R^2^ = 0.83±0.012, VAF = 0.89±0.005), but decreased in the largest perturbation magnitude (R^2^ = 0.64±0.039, VAF = 0.84±0.048). The inclusion of sensorimotor β activity as a predictor of balance-correcting muscle activity also allowed the hSRM to reconstruct the second burst of balance-correcting muscle activity (*Figure 9A*). The inclusion of the longer latency feedback loop in the hSRM resulted in more accurate reconstructions of muscle activity across all perturbation magnitudes compared to the single-loop mSRM (R^2^: t(74) = -2.892, p = 0.005; VAF: t(74) = -3.499, p < 0.001). To further compare these models, we performed an AIC analysis and found that the hSRM when using sensorimotor β activity as a predictor (AIC = -353958) was a better model of perturbation evoked muscle activity than the single-loop mSRM (AIC = -352106), but worse than using either cortical N1 or CoM kinematics as a predictor of longer latency muscle activity. The cortical feedback gain associated with sensorimotor β’s contribution to balance-correcting muscle activity (k_β_) significantly increased with perturbation magnitude (F(1.89,24.52) = 7.77, p = 0.003) (*Figure 9B*).

## Discussion

Our results provide experimental and computational evidence supporting the progressive recruitment of cortical resources in balance control as task difficulty increases. Our data support the hypothesis that in addition to subcortically-mediated balance-correcting muscle activity, cortically-mediated muscle activity is recruited on an individual basis as balance challenge increased at latencies consistent with longer transcortical sensorimotor loops. We further show that balance-correcting muscle activity can be explained via subcortical and cortical sensorimotor feedback loops that process similar sensory information–particularly CoM kinematic error signals–at different intrinsic neural delays. Cortical activity during reactive balance recovery further support the role of transcortical circuits in generating later balance-correcting muscle activity. Our study thus provides a computational framework for dissociating cortical and subcortical muscle activity without need for direct recordings from the brain. Quantifying cortical contributions to balance may be important for assessing balance ability in health and disease, as well as to optimize training, rehabilitation, and assistive devices.

Our results support the hypothesis that long-latency responses to perturbations consist of an initial subcortically-mediated response, followed by a cortical response that becomes engaged as balance task difficulty increases. In both the upper and lower limbs, muscle responses to an unexpected mechanical perturbation follow a stereotypical sequence of short-, and long-latency responses [32,32–34,61]. The spinally-mediated short-latency responses are largely absent in reactive balance recovery in the support surface translations used here [14–17,24,37], and are insufficient to recover balance on their own [28]. In contrast, long-latency responses require the brainstem [8,9,12,62], and are capable of restoring balance [14–16,31,33,60,63–65]. Two components of long-latency muscle activity (LLR1, LLR2) have been identified in both the upper [32,34,61] and lower limb [33,34], with the first being more brainstem mediated (LLR1) and the second being more cortically-mediated (LLR2) [11,21,61]. As such we modeled balance-correcting muscle activity by hierarchical sensorimotor feedback circuits processing balance-error signals at two different loop delays and thresholds.

Here we provide the first evidence that the later cortically-mediated burst of muscle activity (LLR2) appears at individual-specific difficulty levels, potentially once the earlier subcortically-mediated activity (LLR1) becomes inadequate. The first burst was present at all perturbation magnitudes and remained less affected by perturbation magnitude while the second, presumably cortically mediated, burst increased markedly with increasing balance challenge. While we have not previously seen these second bursts consistently in young adults, our prior study of older adults and individuals with Parkinson’s disease frequently exhibited second bursts that could not be accounted for by a single sensorimotor loop [17]. Our revised neuromechanical model explicitly tested the hypothesis that brainstem and cortical circuits process similar sensory information, resulting in two sequential bursts of muscle activity that are both shaped by CoM acceleration signals. In contrast, our prior model with a single delay was insufficient to reproduce muscle activity, particularly at the largest perturbation levels. Here we show that the second burst of activity is primarily driven by CoM acceleration feedback, similarly to the initial burst of muscle activity [31] further corroborating that similar sensorimotor transformations for balance control may occur in parallel at the cortical and subcortical levels. It remains to be seen whether the model also explains the second burst of muscle activity seen in older adults with and without Parkinson’s disease who are at higher risk for falls [17].

In contrast to our prior studies of this same dataset [24,25], we did not find scaling of N1 or β activity with perturbation magnitude in the data analyzed here. The lack of scaling of cortical activity could have arisen from the more selected data set used in this study, which excluded trials in which participants took a step thereby excluding 2 participants from analysis at the largest perturbation magnitude. Additionally, some participants included in previous studies were excluded from this analysis due to a lack of kinematic or EMG data, preventing SRM analysis. Furthermore, the lack of scaling of N1 and β power with perturbation magnitude compared to our prior study may also have resulted from the more rigorous EEG preprocessing pipeline used here to remove muscle and motion artifacts. We also had smaller N1 amplitudes than those reported previously because of the average referencing method used here vs. mastoid electrode reference (TP9/TP10) used previously [25]. The lower β power observed here could be due to the removal of muscle artifacts that include substantial power in the β range [66–68].

The timing and temporal features of cortical activity evoked during balance perturbations are consistent with their potential role in driving the second burst of muscle activity. Perturbation evoked cortical N1 and β activity are observed at a similar latency as the initial burst of balance correcting muscle activity, thus these signals cannot drive the initial, presumed subcortical, burst of muscle activity (LLR1). The N1 is larger on trials requiring stepping response [37,69], however, it has yet to be demonstrated whether cortical N1 drives this subsequent motor behavior. Our results show that similar to muscle activity, the cortical N1 and β activity may also be driven by CoM acceleration signals, and occur early enough to drive the second, presumed cortically-mediated burst of muscle activity (LLR2). Nevertheless, we always observed perturbation evoked cortical N1 and an increase β power, even at the lowest perturbation magnitude when the second, presumed cortically-mediated burst was often absent. Therefore, the cortex may receive ascending sensory information encoding balance error but may not always generate a balance-correcting muscle response until the level of balance challenge is sufficiently high. This coupled with the lack of increase in EEG activity between perturbation magnitudes may suggest that these cortical signatures play a primary role in detecting balance error, which could initiate a separate mechanism that gates *descending* cortical contributions to balance correcting muscle activity.

Our findings align with the concept of the N1 acting as an “alarm signal” that triggers downstream effects that may influence behavior, similar to the proposed role of the cognitive error-related negativity (ERN) [70,71]. The perturbation evoked cortical N1 is thought to be an error assessment signal [36,72] as it exhibits increased amplitude in conditions of heightened threat [73] and is absent when perturbations are predictable [39]. Additionally, N1 tends to be larger in individuals with worse balance [25]. These findings collectively suggest that the balance N1 is not solely linked to sensory integration but likely serves as an indicator of error detection and may share a common underlying mechanism with cognitive ERN [36,72]. However, the perturbation evoked cortical N1 may be more capable of testing relationships between error detection and subsequent behavior than the cognitive ERN as it occurs in a more naturalistic behavior with potentially severe consequences associated with task failure [71]. The cortical N1 has been localized to the supplementary motor area (SMA) [69,74,75] making it a prime contender to drive subsequent motor activity, as the SMA has both afferent and efferent connections to the primary motor cortex [76], prefrontal cortex [76,77] as well as connections with brainstem structures [78]. SMA plays a crucial role in action monitoring and may initiate downstream activity in other neural structures [79] and is heavily implicated in self-initiated movements [80], motor planning and execution [81,82], response inhibition [83], and action sequencing [81,82,84]. Therefore, it is conceivable that N1 may initiate a cascading effect leading to cortical contributions to reactive balance, given that N1 is consistently observed across all magnitudes, whereas cortical contributions to balance-correcting muscle activity are not present until the level of balance challenge is sufficiently high.

Our hierarchical neuromechanical model may be useful in rehabilitation applications due to its ability to quantify increased cortical engagement in balance control without need for direct measurement of brain activity. Our ability to decompose balance correcting muscle activity into subcortical and cortical components may allow for our hSRM model to be used as a tool to index shifts in hierarchical motor control. Evidence suggests that as balance health declines, balance control progressively shifts to be more cortically mediated [11,18–23]. However, the engagement of cortical resources is typically assessed behaviorally, or with cortical blood flow measures [85] that follow neural activity, with both focused primarily on prefrontal contributions to balance. In contrast, assessment of the second burst of muscle activity may provide an index of the cortical contributions to balance control that occur prior to significant prefrontal engagement [45].

Following this logic, it is feasible that the availability of both cortical and subcortical resources have already been compromised by the time of prefrontal engagement, and/or an individual’s first fall [86–89]. Reactive balance responses could therefore be used to assess individual differences in cortical control of balance, and to tune the level of difficulty to optimize rehabilitation training [90,91], or possibly even tune assistive devices.

## References

1. Bard P, Macht MB. The Behaviour of Chronically Decerebrate Cats. Ciba Foundation Symposium - Neurological Basis of Behaviour. John Wiley & Sons, Ltd; 1958. pp. 55–75. doi:10.1002/9780470719091.ch4

2. Lawrence DG, Kuypers HGJM. THE FUNCTIONAL ORGANIZATION OF THE MOTOR SYSTEM IN THE MONKEY1: II. THE EFFECTS OF LESIONS OF THE DESCENDING BRAIN-STEM PATHWAYS. Brain. 1968;91: 15–36. doi:10.1093/brain/91.1.15

3. Beloozerova IN, Sirota MG. The role of the motor cortex in the control of accuracy of locomotor movements in the cat. The Journal of Physiology. 1993;461: 1–25. doi:10.1113/jphysiol.1993.sp019498

4. Beloozerova IN, Sirota MG, Swadlow HA, Orlovsky GN, Popova LB, Deliagina TG. Activity of Different Classes of Neurons of the Motor Cortex during Postural Corrections. J Neurosci. 2003;23: 7844–7853. doi:10.1523/JNEUROSCI.23-21-07844.2003

5. Beloozerova IN, Sirota MG, Swadlow HA. Activity of Different Classes of Neurons of the Motor Cortex during Locomotion. J Neurosci. 2003;23: 1087–1097. doi:10.1523/JNEUROSCI.23-03-01087.2003

6. Drew T, Prentice S, Schepens B. Cortical and brainstem control of locomotion. Progress in Brain Research. 2004;143: 251–261.

7. Drew T, Andujar J-E, Lajoie K, Yakovenko S. Cortical mechanisms involved in visuomotor coordination during precision walking. Brain Research Reviews. 2008;57: 199–211. doi:10.1016/j.brainresrev.2007.07.017

8. Deliagina TG, Orlovsky GN, Zelenin PV, Beloozerova IN. Neural Bases of Postural Control. Physiology. 2006;21: 216–225. doi:10.1152/physiol.00001.2006

9. Deliagina TG, Zelenin PV, Beloozerova IN, Orlovsky GN. Nervous mechanisms controlling body posture. Physiology & Behavior. 2007;92: 148–154. doi:10.1016/j.physbeh.2007.05.023

10. Deliagina TG, Beloozerova IN, Zelenin PV, Orlovsky GN. Spinal and supraspinal postural networks. Brain Research Reviews. 2008;57: 212–221. doi:10.1016/j.brainresrev.2007.06.017

11. Maki BE, McIlroy WE. Cognitive demands and cortical control of human balance-recovery reactions. J Neural Transm. 2007;114: 1279–1296. doi:10.1007/s00702-007-0764-y

12. Honeycutt CF, Gottschall JS, Nichols TR. Electromyographic Responses From the Hindlimb Muscles of the Decerebrate Cat to Horizontal Support Surface Perturbations. Journal of Neurophysiology. 2009;101: 2751–2761. doi:10.1152/jn.91040.2008

13. Clark DJ. Automaticity of walking: functional significance, mechanisms, measurement and rehabilitation strategies. Front Hum Neurosci. 2015;9. doi:10.3389/fnhum.2015.00246

14. Welch TDJ, Ting LH. A Feedback Model Reproduces Muscle Activity During Human Postural Responses to Support-Surface Translations. Journal of Neurophysiology. 2008;99: 1032–1038. doi:10.1152/jn.01110.2007

15. Welch TDJ, Ting LH. A Feedback Model Explains the Differential Scaling of Human Postural Responses to Perturbation Acceleration and Velocity. Journal of Neurophysiology. 2009;101: 3294–3309. doi:10.1152/jn.90775.2008

16. Welch TDJ, Ting LH. Mechanisms of Motor Adaptation in Reactive Balance Control. PLOS ONE. 2014;9: e96440. doi:10.1371/journal.pone.0096440

17. McKay JL, Lang KC, Bong SM, Hackney ME, Factor SA, Ting LH. Abnormal center of mass feedback responses during balance: A potential biomarker of falls in Parkinson’s disease. PLoS One. 2021;16: e0252119. doi:10.1371/journal.pone.0252119

18. Shumway-Cook A, Woollacott M, Kerns KA, Baldwin M. The Effects of Two Types of Cognitive Tasks on Postural Stability in Older Adults With and Without a History of Falls. J Gerontol A Biol Sci Med Sci. 1997;52A: M232–M240. doi:10.1093/gerona/52A.4.M232

19. Rankin JK, Woollacott MH, Shumway-Cook A, Brown LA. Cognitive Influence on Postural StabilityA Neuromuscular Analysis in Young and Older Adults. J Gerontol A Biol Sci Med Sci. 2000;55: M112–M119. doi:10.1093/gerona/55.3.M112

20. Woollacott M, Shumway-Cook A. Attention and the control of posture and gait: a review of an emerging area of research. Gait Posture. 2002;16: 1–14.

21. Jacobs JV, Horak FB. Cortical control of postural responses. Journal of Neural Transmission. 2007;114: 1339. doi:10.1007/s00702-007-0657-0

22. Ozdemir RA, Contreras-Vidal JL, Lee B-C, Paloski WH. Cortical activity modulations underlying agerelated performance differences during posture–cognition dual tasking. Exp Brain Res. 2016;234: 3321–3334. doi:10.1007/s00221-016-4730-5

23. Ozdemir RA, Contreras-Vidal JL, Paloski WH. Cortical control of upright stance in elderly. Mechanisms of Ageing and Development. 2018;169: 19–31. doi:10.1016/j.mad.2017.12.004

24. Ghosn NJ, Palmer JA, Borich MR, Ting LH, Payne AM. Cortical Beta Oscillatory Activity Evoked during Reactive Balance Recovery Scales with Perturbation Difficulty and Individual Balance Ability. Brain Sci. 2020;10. doi:10.3390/brainsci10110860

25. Payne AM, Ting LH. Worse balance is associated with larger perturbation-evoked cortical responses in healthy young adults. Gait & Posture. 2020;80: 324–330. doi:10.1016/j.gaitpost.2020.06.018

26. Horak FB. Adaptation of automatic postural responses. The acquisition of motor behavior in vertebrates. 1996; 57–85.

27. Horak FB. Postural orientation and equilibrium: what do we need to know about neural control of balance to prevent falls? Age and Ageing. 2006;35: ii7–ii11. doi:10.1093/ageing/afl077

28. Horak FB, Macpherson JM. Postural Orientation and Equilibrium. Comprehensive Physiology. American Cancer Society; 2011. pp. 255–292. doi:10.1002/cphy.cp120107

29. Dunbar DC, Horak FB, Macpherson JM, Rushmer DS. Neural control of quadrupedal and bipedal stance: Implications for the evolution of erect posture. American Journal of Physical Anthropology. 1986;69: 93–105. doi:10.1002/ajpa.1330690111

30. Macpherson JM, Fung J. Weight Support and Balance During Perturbed Stance in the Chronic Spinal Cat. Journal of Neurophysiology. 1999;82: 3066–3081. doi:10.1152/jn.1999.82.6.3066

31. Lockhart DB, Ting LH. Optimal sensorimotor transformations for balance. Nature Neuroscience. 2007;10: 1329–1336. doi:10.1038/nn1986

32. Lee RG, Tatton WG. Motor Responses to Sudden Limb Displacements in Primates with Specific CNS Lesions and in Human Patients with Motor System Disorders. Canadian Journal of Neurological Sciences. 1975;2: 285–293. doi:10.1017/S0317167100020382

33. Taube W, Schubert M, Gruber M, Beck S, Faist M, Gollhofer A. Direct corticospinal pathways contribute to neuromuscular control of perturbed stance. J Appl Physiol. 2006;101: 420–429. doi:10.1152/japplphysiol.01447.2005

34. Pruszynski JA, Scott SH. Optimal feedback control and the long-latency stretch response. Exp Brain Res. 2012;218: 341–359. doi:10.1007/s00221-012-3041-8

35. Payne AM, Hajcak G, Ting LH. Dissociation of muscle and cortical response scaling to balance perturbation acceleration. Journal of Neurophysiology. 2019;121: 867–880. doi:10.1152/jn.00237.2018

36. Payne AM, Ting LH, Hajcak G. Do sensorimotor perturbations to standing balance elicit an errorrelated negativity? Psychophysiology. 2019;56: e13359. doi:10.1111/psyp.13359

37. Payne AM, Ting LH. Balance perturbation-evoked cortical N1 responses are larger when stepping and not influenced by motor planning. Journal of Neurophysiology. 2020;124: 1875–1884. doi:10.1152/jn.00341.2020

38. Quant S, Adkin AL, Staines WR, McIlroy WE. Cortical activation following a balance disturbance. Exp Brain Res. 2004;155: 393–400. doi:10.1007/s00221-003-1744-6

39. Adkin AL, Quant S, Maki BE, McIlroy WE. Cortical responses associated with predictable and unpredictable compensatory balance reactions. Exp Brain Res. 2006;172: 85. doi:10.1007/s00221-005-0310-9

40. Peterson SM, Ferris DP. Differentiation in Theta and Beta Electrocortical Activity between Visual and Physical Perturbations to Walking and Standing Balance. eNeuro. 2018;5. doi:10.1523/ENEURO.0207-18.2018

41. Peterson SM, Ferris DP. Group-level cortical and muscular connectivity during perturbations to walking and standing balance. Neuroimage. 2019;198: 93–103. doi:10.1016/j.neuroimage.2019.05.038

42. Dietz V, Quintern J, Berger W. Cerebral evoked potentials associated with the compensatory reactions following stance and gait perturbation. Neuroscience Letters. 1984;50: 181–186. doi:10.1016/0304-3940(84)90483-X

43. Mierau A, Pester B, Hülsdünker T, Schiecke K, Strüder HK, Witte H. Cortical Correlates of Human Balance Control. Brain Topogr. 2017;30: 434–446. doi:10.1007/s10548-017-0567-x

44. Solis-Escalante T, van der Cruijsen J, de Kam D, van Kordelaar J, Weerdesteyn V, Schouten AC. Cortical dynamics during preparation and execution of reactive balance responses with distinct postural demands. Neuroimage. 2019;188: 557–571. doi:10.1016/j.neuroimage.2018.12.045

45. Palmer JA, Payne AM, Ting LH, Borich MR. Cortical Engagement Metrics During Reactive Balance Are Associated With Distinct Aspects of Balance Behavior in Older Adults. Front Aging Neurosci. 2021;0. doi:10.3389/fnagi.2021.684743

46. Salenius S, Schnitzler A, Salmelin R, Jousmäki V, Hari R. Modulation of human cortical rolandic rhythms during natural sensorimotor tasks. Neuroimage. 1997;5: 221–228. doi:10.1006/nimg.1997.0261

47. Gilbertson T, Lalo E, Doyle L, Lazzaro VD, Cioni B, Brown P. Existing Motor State Is Favored at the Expense of New Movement during 13-35 Hz Oscillatory Synchrony in the Human Corticospinal System. J Neurosci. 2005;25: 7771–7779. doi:10.1523/JNEUROSCI.1762-05.2005

48. Lalo E, Gilbertson T, Doyle L, Lazzaro VD, Cioni B, Brown P. Phasic increases in cortical beta activity are associated with alterations in sensory processing in the human. Exp Brain Res. 2007;177: 137–145. doi:10.1007/s00221-006-0655-8

49. Pogosyan A, Gaynor LD, Eusebio A, Brown P. Boosting Cortical Activity at Beta-Band Frequencies Slows Movement in Humans. Current Biology. 2009;19: 1637–1641. doi:10.1016/j.cub.2009.07.074

50. Barone J, Rossiter HE. Understanding the Role of Sensorimotor Beta Oscillations. Front Syst Neurosci. 2021;15: 655886. doi:10.3389/fnsys.2021.655886

51. Engel AK, Fries P. Beta-band oscillations—signalling the status quo? Current Opinion in Neurobiology. 2010;20: 156–165. doi:10.1016/j.conb.2010.02.015

52. van Wijk BCM, Beek PJ, Daffertshofer A. Neural synchrony within the motor system: what have we learned so far? Front Hum Neurosci. 2012;0. doi:10.3389/fnhum.2012.00252

53. Kilavik BE, Zaepffel M, Brovelli A, MacKay WA, Riehle A. The ups and downs of beta oscillations in sensorimotor cortex. Experimental Neurology. 2013;245: 15–26. doi:10.1016/j.expneurol.2012.09.014

54. Delorme A, Makeig S. EEGLAB: an open source toolbox for analysis of single-trial EEG dynamics including independent component analysis. Journal of Neuroscience Methods. 2004;134: 9–21. doi:10.1016/j.jneumeth.2003.10.009

55. de Cheveigné A. ZapLine: A simple and effective method to remove power line artifacts. NeuroImage. 2020;207: 116356. doi:10.1016/j.neuroimage.2019.116356

56. Klug M, Kloosterman NA. Zapline-plus: A Zapline extension for automatic and adaptive removal of frequency-specific noise artifacts in M/EEG. Human Brain Mapping. 2022;43: 2743–2758. doi:10.1002/hbm.25832

57. Palmer JA, Kreutz-Delgado K, Makeig S. AMICA: An Adaptive Mixture of Independent Component Analyzers with Shared Components. 2011; 15.

58. Makeig S. Auditory event-related dynamics of the EEG spectrum and effects of exposure to tones. Electroencephalography & Clinical Neurophysiology. 1993;86: 283–293. doi:10.1016/0013-4694(93)90110-H

59. Basmajian JV. Electrode placement in EMG biofeedback. Baltimore: Williams & Wilkins; 1980.

60. Safavynia SA, Ting LH. Long-latency muscle activity reflects continuous, delayed sensorimotor feedback of task-level and not joint-level error. J Neurophysiol. 2013;110: 1278–1290. doi:10.1152/jn.00609.2012

61. Pruszynski JA, Kurtzer I, Scott SH. The long-latency reflex is composed of at least two functionally independent processes. J Neurophysiol. 2011;106: 449–459. doi:10.1152/jn.01052.2010

62. Takakusaki K, Saitoh K, Harada H, Kashiwayanagi M. Role of basal ganglia-brainstem pathways in the control of motor behaviors. Neurosci Res. 2004;50: 137–151. doi:10.1016/j.neures.2004.06.015

63. Nashner LM. Fixed patterns of rapid postural responses among leg muscles during stance. Exp Brain Res. 1977;30: 13–24. doi:10.1007/BF00237855

64. Ting LH, Macpherson JM. Ratio of shear to load ground-reaction force may underlie the directional tuning of the automatic postural response to rotation and translation. J Neurophysiol. 2004;92: 808–823. doi:10.1152/jn.00773.2003

65. Ting LH. Dimensional reduction in sensorimotor systems: a framework for understanding muscle coordination of posture. Cisek P, Drew T, Kalaska JF, editors. Progress in Brain Research. 2007;165: 299–321. doi:10.1016/S0079-6123(06)65019-X

66. Muthukumaraswamy S. High-frequency brain activity and muscle artifacts in MEG/EEG: A review and recommendations. Frontiers in Human Neuroscience. 2013;7. Available: https://www.frontiersin.org/articles/10.3389/fnhum.2013.00138

67. Chaumon M, Bishop DVM, Busch NA. A practical guide to the selection of independent components of the electroencephalogram for artifact correction. Journal of Neuroscience Methods. 2015;250: 47–63. doi:10.1016/j.jneumeth.2015.02.025

68. Chen X, Xu X, Liu A, Lee S, Chen X, Zhang X, et al. Removal of Muscle Artifacts From the EEG: A Review and Recommendations. IEEE Sensors Journal. 2019;19: 5353–5368. doi:10.1109/JSEN.2019.2906572

69. Solis-Escalante T, Stokkermans M, Cohen MX, Weerdesteyn V. Cortical responses to whole-body balance perturbations index perturbation magnitude and predict reactive stepping behavior. European Journal of Neuroscience. 2020;n/a. 10.1111/ejn.14972

70. Cavanagh JF, Frank MJ. Frontal theta as a mechanism for cognitive control. Trends in Cognitive Sciences. 2014;18: 414–421. doi:10.1016/j.tics.2014.04.012

71. LoTemplio SB, Lopes CL, McDonnell AS, Scott EE, Payne BR, Strayer DL. Updating the relationship of the Ne/ERN to task-related behavior: A brief review and suggestions for future research. Frontiers in Human Neuroscience. 2023;17. Available: https://www.frontiersin.org/articles/10.3389/fnhum.2023.1150244

72. Payne AM, Ting LH, Hajcak G. The balance N1 and the ERN correlate in amplitude across individuals in small samples of younger and older adults. Exp Brain Res. 2023 [cited 12 Sep 2023]. doi:10.1007/s00221-023-06692-9

73. Adkin AL, Campbell AD, Chua R, Carpenter MG. The influence of postural threat on the cortical response to unpredictable and predictable postural perturbations. Neuroscience Letters. 2008;435: 120–125. doi:10.1016/j.neulet.2008.02.018

74. Marlin A, Mochizuki G, Staines WR, McIlroy WE. Localizing evoked cortical activity associated with balance reactions: does the anterior cingulate play a role? Journal of Neurophysiology. 2014;111: 2634–2643. doi:10.1152/jn.00511.2013

75. Mierau A, Hülsdünker T, Strüder HK. Changes in cortical activity associated with adaptive behavior during repeated balance perturbation of unpredictable timing. Front Behav Neurosci. 2015;9. doi:10.3389/fnbeh.2015.00272

76. Jürgens U. The efferent and afferent connections of the supplementary motor area. Brain Res. 1984;300: 63–81. doi:10.1016/0006-8993(84)91341-6

77. Hertrich I, Dietrich S, Ackermann H. The role of the supplementary motor area for speech and language processing. Neuroscience & Biobehavioral Reviews. 2016;68: 602–610. doi:10.1016/j.neubiorev.2016.06.030

78. Çavdar S, Köse B, Altınöz D, Özkan M, Günes YC, Algin O. The brainstem connections of the supplementary motor area and its relations to the corticospinal tract: Experimental rat and human 3-tesla tractography study. Neuroscience Letters. 2023;798: 137099. doi:10.1016/j.neulet.2023.137099

79. Bonini F, Burle B, Liégeois-Chauvel C, Régis J, Chauvel P, Vidal F. Action Monitoring and Medial Frontal Cortex: Leading Role of Supplementary Motor Area. Science. 2014;343: 888–891. doi:10.1126/science.1247412

80. Passingham RE, Bengtsson SL, Lau HC. Medial frontal cortex: from self-generated action to reflection on one’s own performance. Trends in Cognitive Sciences. 2010;14: 16–21. doi:10.1016/j.tics.2009.11.001

81. Tanji J. New concepts of the supplementary motor area. Current Opinion in Neurobiology. 1996;6: 782–787. doi:10.1016/S0959-4388(96)80028-6

82. Nachev P, Kennard C, Husain M. Functional role of the supplementary and pre-supplementary motor areas. Nat Rev Neurosci. 2008;9: 856–869. doi:10.1038/nrn2478

83. Mostofsky SH, Simmonds DJ. Response Inhibition and Response Selection: Two Sides of the Same Coin. Journal of Cognitive Neuroscience. 2008;20: 751–761. doi:10.1162/jocn.2008.20500

84. Tanji J. Sequential organization of multiple movements: involvement of cortical motor areas. Annu Rev Neurosci. 2001;24: 631–651. doi:10.1146/annurev.neuro.24.1.631

85. Stuart S, Vitorio R, Morris R, Martini DN, Fino PC, Mancini M. Cortical activity during walking and balance tasks in older adults and Parkinson’s disease: a structured review. Maturitas. 2018;113: 53–72. doi:10.1016/j.maturitas.2018.04.011

86. Reuter-Lorenz PA, Cappell KA. Neurocognitive Aging and the Compensation Hypothesis. Curr Dir Psychol Sci. 2008;17: 177–182. doi:10.1111/j.1467-8721.2008.00570.x

87. Chatterjee SA, Fox EJ, Daly JJ, Rose DK, Wu SS, Christou EA, et al. Interpreting Prefrontal Recruitment During Walking After Stroke: Influence of Individual Differences in Mobility and Cognitive Function. Front Hum Neurosci. 2019;13: 194. doi:10.3389/fnhum.2019.00194

88. Clark DJ, Manini TM, Ferris DP, Hass CJ, Brumback BA, Cruz-Almeida Y, et al. Multimodal Imaging of Brain Activity to Investigate Walking and Mobility Decline in Older Adults (Mind in Motion Study): Hypothesis, Theory, and Methods. Front Aging Neurosci. 2020;11. doi:10.3389/fnagi.2019.00358

89. St George RJ, Hinder MR, Puri R, Walker E, Callisaya ML. Functional Near-infrared Spectroscopy Reveals the Compensatory Potential of Pre-frontal Cortical Activity for Standing Balance in Young and Older Adults. Neuroscience. 2021;452: 208–218. doi:10.1016/j.neuroscience.2020.10.027

90. Guadagnoli MA, Lee TD. Challenge Point: A Framework for Conceptualizing the Effects of Various Practice Conditions in Motor Learning. Journal of Motor Behavior. 2004;36: 212–224. doi:10.3200/JMBR.36.2.212-224

91. Sawers A, Allen JL, Ting LH. Long-term training modifies the modular structure and organization of walking balance control. J Neurophysiol. 2015;114: 3359–3373. doi:10.1152/jn.00758.2015

